# Formalization of Genome Interval Relations

**DOI:** 10.1101/006650

**Authors:** Chris J. Mungall

**Affiliations:** Genomics Division, Lawrence Berkeley National Laboratory, 1 Cycltotron Road MS 64-121, Berkeley, CA 94720, USA

## Abstract

In order to take full advantage of next generation genomics data, I need informatics methods to be based on agreed upon *formally specified* standards that can be implemented easily in a uniform fashion without ambiguity. These standards should be encoded as logical formulae, so that provably correct and efficient decision procedures can be used for query answering and validation.

In this paper I present the core of such a standard for sequence data: a collection of definitions of relations that hold between genomic intervals, and an alegbra for performing operations upon these intervals. I show how these relations can be used to extend formalize concepts in the Sequence Ontology (SO).

## 1 Introduction

### 1.1 Genome Databases and the need for inference

The genome of every organism is organized as a collection of sequentially ordered nucleotide bases, with distinct regions of the sequence comprising a variety of structural and functional elements: genes, exons, regulatory regions, untranslated regions and so on. A precise understanding of these elements is vital for the life sciences and clinical applications, yet despite this importance, there is no formally defined terminology for describing the different relations that can hold at the primary structure level. The lack of any such standard is a hindrance to interoperability between computer systems, which could have serious ramifications as genomic database continue to grow in size and importance.

To illustrate the problem, consider basic biological questions such as *how many distinct introns are there in the human genome?, what is the proportion of size of intergenic to genic regions across all sequenced eukaryotes?* and *what is the ratio of SNPs in coding regions vs those in UTRs?* These are reasonable questions that turn out to be difficult to answer without recourse to the suboptimal solution of writing programs to obtain the answers. One reason existing databases have problems answering these questions is that they are each inconsistent in what information is represented and what information must be derived algorithmically. A collection of formal genome interval relations would be of use not only for precisely specifying region-based queries, but also as the basis for computable definitions of the inference rules required to obtain a set of introns given a set of exons, or to obtain UTRs and coding sequence from transcript and start and stop codon information.

These relations should extend existing work on defining relations in biology[1], in which a formal or semi-formal approach is adopted, with the definitions of relations specified as unambiguously as possible. One way to eliminate ambiguity is to specify relations using First-Order Logic (FOL) axioms. This allows the use of theorem provers to determine the consequences of the statements.

### 1.2 Sequence Ontology

The Sequence Ontology (SO)[2] was originally conceived of as a structured controlled vocabulary for genome databases and exchange formats. The SO can also be treated as a collection of axioms stating truth-conditions for relations between instances of biological features such as genes, transcripts and exons. The translation is between SO relationships and FOL axioms is shown in table 1.

**Table 1:** Translation table for relationships in the Sequence Ontology (SO). The convention of using italics for type level relations and bold for instance level relations is taken from[6]. The rules for translating to instances from a genome database are not specified.

| Relationship | Axiom |
| --- | --- |
| $\langle T \text{ is\_a } U \rangle$ | $T(x) \rightarrow U(x)$ |
| $\langle T R U \rangle$ | $T(x) \rightarrow \exists y : U(y), R(x, y)$ |
| $\langle T \text{ is\_a } G \text{ that } D \rangle$ | $T(x) \leftrightarrow G(x), \forall \langle R, U \rangle \in D : (\exists y : U(y), R(x, y))$ |
| $\langle \text{transitive } R \rangle$ | $R(x, y), R(y, z) \rightarrow R(x, z)$ |

We can translate each feature type *T* in SO to a unary predicate such that *T*(*x*) is true whenever *x* is an instance of *T*. This exon(*x*) holds for all values of x where x is an instance of an exon. Here we treat SO as a representation of what the Basic Formal Ontology calls *generically dependent continuants*[3]. This means that a single instance of a SO type can have multiple molecular bearers: an individual human consists in part of trillions of chromosome molecule instances which (excluding somatic variation) in toto bear a single genome instance. It is these instances that form the domain of discourse of genome databases, rather than the individual chromosome molecules. We can derive fact axioms such as `exon(ENSE00001545001)` from rows in a relational database such as Ensembl[4] or Chado[5].

The SO *is_a* hierarchy is translated such that 〈*T is_a U*〉 becomes *T*(*x*) → *U*(*x*). For example `mRNA`(*x*) → `transcript`(*x*) (every mRNA is a transcript). All-some relationships such as *part_of* specified using the methodology of the Relations Ontology[6] can be translated to quantified statements over individuals, such that for example the SO type-level relationship 〈`TSS` *part_of* `transcript`〉 becomes `TSS`(*x*) → ∃*y*, `transcript`(*y*), `part_of`(*x*, *y*). SO also includes relation axioms, such as transitivity of the *part_of* relation.

Compositional terms in SO generally have logical definitions, stating necessary and sufficient conditions expressed in simple genus differentia form, i.e. *an X is a G that D*, for example, 〈`transposable_element_gene` *is_a* gene that is *part_of* a `transposable_element`〉. This translates to the definitional axiom:

`transposable_element_gene`(*x*) ↔

`gene`(*x*) ∧ (∃*y*: `transposable_element`(*y*), `part_of`(*x*, *y*))

These logical definitions can be used by reasoning engines to automatically classify the ontology (i.e. infer *is_a* relations). They can also be translated to relational database queries to find instances of implicit feature types.

`SO` is currently lacking computable logical definitions for many terms such as `UTR`, `region`, `five_prime_UTR`, `CDS`, `intron` and `five_prime_coding_exon`. These definitions are currently specified in natural language, which means they must be translated by humans if they are to be used algorithmically (e.g. to query a database for all implicit `intron` or `five_prime_UTR` features by performing arithmetic operations on the ranges of stated features). SO is also lacking axioms constraining the genomic positioning of related features. For example, that each `exon` must lie within the region of the gene of which it is a part, or that the `TSS` of a `transcript` must be upstream of that transcript.

Ideally we would like to add these kinds of logical definitions and axioms to SO, but we first need to extend on the set of relations used in SO. We can build upon existing relations designed to support qualitative spatial and temporal reasoning, namely the Region Connection Calculus and the Allen Interval Algebra.

### 1.3 Region Connection Calculus (RCC-8)

The region connection calculus (RCC) is a system for qualitative spatial representation and reasoning. RCC abstractly describes regions (in Euclidian space, or in a topological space) by their possible relations to each other. RCC8 consists of 8 basic relations that can hold between two regions: *disconnected (DC), externally connected (EC), equal (EQ), partially overlapping (PO), tangential proper part (TPP), tangential proper part inverse (TPPi), non-tangential proper part (NTPP)* and *non-tangential proper part inverse (NTPPi)*. See figure 1.

**Figure 1:**
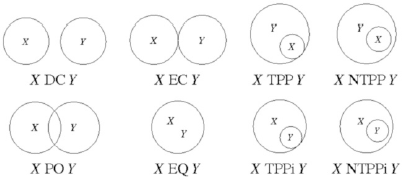
Relations in the Region Connection Calculus

We can compose these relations using logical operators. For example, For example *PP* = *TPP*∪*NTPP*. For qualitative reasoning about the relations between regions, there is a *composition table*[7]. Given the relation *R*_1_ between *x* and *y* and the relation *R*_2_ between *y* and *z*, the composition table allows us to determine *R*_3_, the relation between *x* and *z*. We write this as *R*_1_ · *R*_2_ → *R*_3_, which translates to *xR*_1_*y*, *yR*_2_*z* → *xR*_3_*z*. For example *NTPP* · *NTPP* → *NTPP* (i.e transitivity of non-tangential proper part).

RCC-8 is commonly used in geographical information systems. It could also be of use in reasoning about biological or biochemical entities in any number of dimensions. One consideration is that RCC-8 operates over continuous regions, rather than discrete units, such as nucleotides.

### 1.4 Allen’s Interval Algebra

Allen’s Interval Algebra (AIA)[8] defines possible relations between time intervals, and operations on these intervals, that can be used as a basis for qualitative reasoning about temporal descriptions of events.

The AIA defines 13 pairwise disjoint base relations that capture all possible relations between two intervals. The 13 relations consist of 6 asymmetric relations: *precedes (p), meets (m), overlaps (o), finishes (f), contains (di), starts (s)*, their inverses: *precededBy (pi), metBy (mi), overlappedBy (oi), finishedBy (fi), containedBy*^1^ *(c), startedBy (si)*, and the symmetric *equals (e)* relation.

Composing relations together gives a total possible 8192 relations. The composition operators are intersection (∩), union (∪) and complementation (¬). For example, the union *p* ∪ *pi* holds whenever two time intervals have no time point in common.

Satisfiability is NP-Complete with AIA i.e. given a collection of intervals and the relations that hold between them we cannot in general compute if there are time values for which the relations are true. However, there are tractable sub-algebras for which efficient decision procedures exist[9].

Whilst the AIA is generally described as consisting of temporal intervals, it can be applied to any kind of interval, including genomic intervals. One consideration is that like RCC-8, the AIA assumes continuous intervals, rather than discrete intervals.

## 2 Results

### 2.1 An Algebra of Genomic Intervals

We base our Genomic Interval Algebra (GIA) on the Allen Interval Algebra (AIA), and extend it with additional relations requires for reasoning about DNA sequences. AIA is more suited than RCC-8 for a basis as genome intervals, like tenporal intervals are directional. We change some of the terminology of Allen (e.g. using “upstream” and “downstream” instead of “precedes” and “precededBy”), and take some terms from RCC-8 (e.g. “adjacent”), but use Allen-based definitions.

When considering the biological meaning of these relations, it is important to stress that they hold between *sequences or sequence intervals* (i.e. primary structures) and not the molecules that are the bearers of those sequences. For example, an RNA molecule or intron may exhibit connectedness/adjacency between bases at the secondary structure level. Similarly, a transcription factor protein may exhibit binding between its amino acid chain and the DNA sequence upstream of a gene. We consider both these cases to be non-adjacent and disconnected at the sequence level.

The core of the GIA consists of 16 basic relations *R*^16^ that can hold between any two intervals on the same strand of a sequence, defined in terms of Allen relations. Whilst the IAI treats intervals as primitives, we also provide definitions in terms of junctions (the equivalent being time-points in a temporal calculus), yielding a junction calculus. We then extend the core set of relations to account for strandedness, deriving an additional 32 relations.

Figure 2 illustrates these relations with a simplified example.

**Figure 2:**
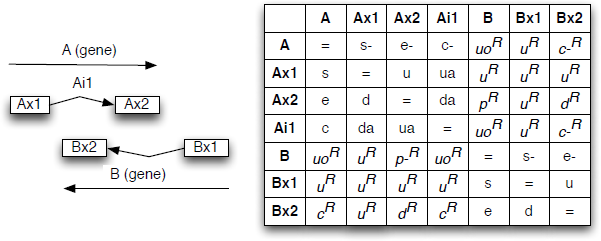
Example of genomic sequence interval relations: two interleaved genes A and B on opposite strands. The lookup table shows the mnemonics for the relations between any two feature intervals. To determine the relation xRy, look up (row:x column:y). For example the first exon of A (Ax1) is upstream of the reverse-complement projection of all the exons of B.

#### 2.1.1 Basic single-strand relations

We have declared 16 interval relations that can hold between two intervals on the same strand of a sequence. These are shown in table 2.

**Table 2:**
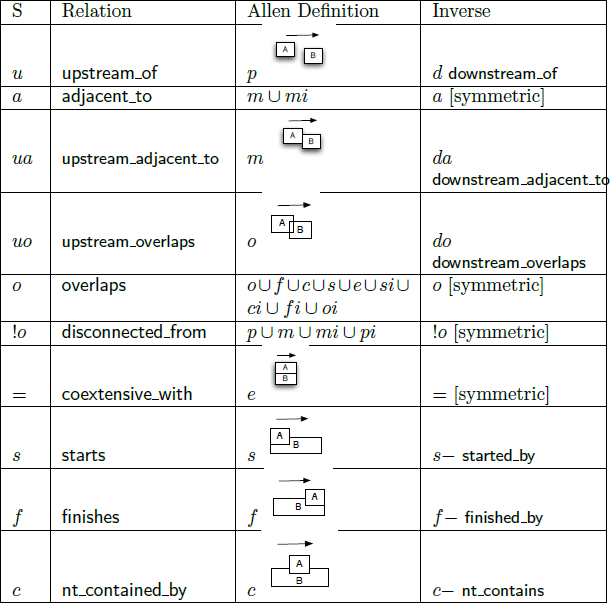
16 Core Relations in the Genome Interval Algebra. Glyphs depict the relation holding between A and B, with the strand indicated by an arrow. This table provides definitions based on Allen relations. We provide both human-friendly names (e.g. `upstream_of`) and mnemonics (e.g. *u*)

| S | Relation | Allen Definition | Inverse |
| --- | --- | --- | --- |
| *u* | upstream_of | $p$ | $d$ downstream_of |
| *a* | adjacent_to | $m \cup mi$ | $a$ [symmetric] |
| *ua* | upstream_adjacent_to | $m$ | $da$<br>downstream_adjacent_to |
| *uo* | upstream_overlaps | $o$ | $do$<br>downstream_overlaps |
| *o* | overlaps | $o \cup f \cup c \cup s \cup e \cup si \cup ci \cup fi \cup oi$ | $o$ [symmetric] |
| *!o* | disconnected_from | $p \cup m \cup mi \cup pi$ | $!o$ [symmetric] |
| $=$ | coextensive_with | $e$ | $=$ [symmetric] |
| *s* | starts | $s$ | $s -$ started_by |
| *f* | finishes | $f$ | $f -$ finished_by |
| *c* | nt_contained_by | $c$ | $c -$ nt_contains |

Some relations are defined in terms of other relations using relation-intersection and relation-union operators. These are defined as follows:

intersection: *x*(*R* ∩ *S*)*y* ↔ *xRy* ∧ *xSy*
union: *x*(*R* ∪ *S*)*y* ↔ *xRy* ∨ *xSy*
complement: *x*(!*R*)*y* ↔ ¬[*xRy*]

Note that we do not take the same set of primitives as Allen; we choose our set based on utility within genome databases and in the SO. This means that there is not an isomorphic correspondence between the GIA and Allen. We provide Allen-based definitions using relation intersection and and union. Unlike Allen, our set is not pairwise disjoint. For example, `adjacent_to = upstream_adjacent_to ∩ downstream_adjacent_to`. We also make different terminological choices from the IAI. For example, we use `overlaps` in a more general sense, in accord with how this term is typically use in bioinformatics, and how it is defined in the Relation Ontology.

#### 2.1.2 Junction-based definitions

We also provide definitions for all interval relations in terms of point-positions or junctions. We define a *proper junction* as a discrete point connecting two nucleotide bases. Junctions are the union of proper junctions and the outermost boundary points of a sequence (this inclusive definition of junction simplifies the axioms).

We define two functions *α* and *ω* each of which maps an interval to a point (junction), correspoding to the start and end of the interval. We introduce a relation succ which holds between two junctions separated by a base, such that the first is at the 5’ end and the second is at the 3’ end. We overload the symbol < as the transitive version of this relation: *x* < *y* ↔ succ(*x*,*y*) ∨ ∃*z*: (succ(*x*, *z*), *z* < *y*. We use > as the inverse of this relation and define <= as the union of < and = and >= as the union of > and =. In the sequence ontology we give these full names such as `before`.

For any interval, the start is before the end. Formally:

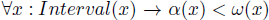

Both < and > are irreflexive, and for any non-circular genome they are anti-symmetric i.e. ¬∃*x*, *y* : *x* < *y*, *y* < *x*.

Table 3 gives the definition of interval relations in terms of junctions.

**Table 3:**
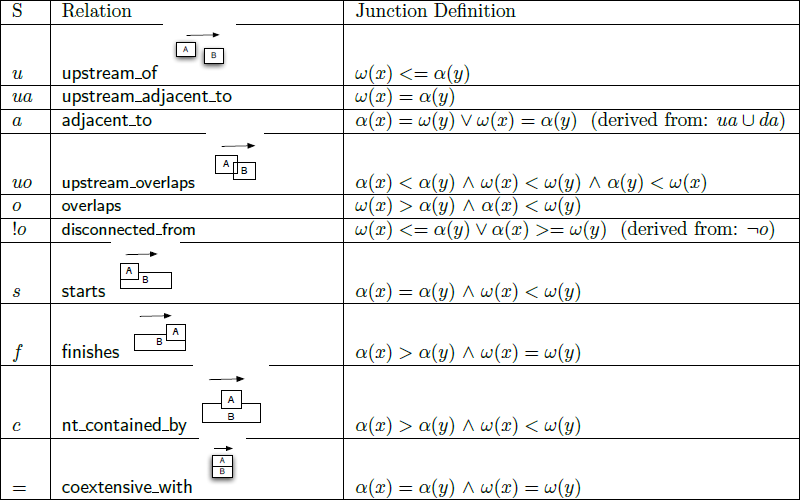
Junction-based definitions. Definitions are not shown for inverses, which can be trivially obtained by reversing *x* and *y*.

| S | Relation | Junction Definition |
| --- | --- | --- |
| $u$ | upstream_of | $\omega(x) \leq \alpha(y)$ |
| $ua$ | upstream_adjacent_to | $\omega(x) = \alpha(y)$ |
| $a$ | adjacent_to | $\alpha(x) = \omega(y) \vee \omega(x) = \alpha(y)$ (derived from: $ua \cup da$) |
| $uo$ | upstream_overlaps | $\alpha(x) < \alpha(y) \wedge \omega(x) < \omega(y) \wedge \alpha(y) < \omega(x)$ |
| $o$ | overlaps | $\omega(x) > \alpha(y) \wedge \alpha(x) < \omega(y)$ |
| $!o$ | disconnected_from | $\omega(x) \leq \alpha(y) \vee \alpha(x) \geq \omega(y)$ (derived from: $\neg o$) |
| $s$ | starts | $\alpha(x) = \alpha(y) \wedge \omega(x) < \omega(y)$ |
| $f$ | finishes | $\alpha(x) > \alpha(y) \wedge \omega(x) = \omega(y)$ |
| $c$ | nt_contained_by | $\alpha(x) > \alpha(y) \wedge \omega(x) < \omega(y)$ |
| $=$ | coextensive_with | $\alpha(x) = \alpha(y) \wedge \omega(x) = \omega(y)$ |

We can eliminate the function terms from the definitions by translating to composition rules such that for example:

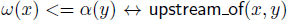

Is translated to:

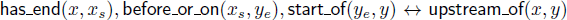

We can eliminate the variables using an equivalence assertion and a relation chain:

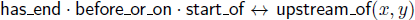

Here `has_start` and `has_end` are a functional relations between a region and a junction, with inverses `start_of` and `end_of`.

These additional relations give us an algebra over junctions and relations which we call *GIA^J^*, which can be used with qualitative reasoning systems. Note the correspondence of succ to the function +1 and between < and > and their arithmetic counterparts - the correspondence allows the use of either artihmetic operations or qualitative reasoning.

#### 2.1.3 Derived Reverse-Complement Relations

The major difference between temporal and genomic intervals is that DNA is stranded. The GIA must therefore be an *extension* of the AIA to fully account for strandedness.

We treat each strand of a double-stranded DNA molecule as bearing two distinct sequence intervals *s*^+^ and *s*^−^, related by the RC relation. Each junction 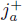 on s^+^ has a unique single cognate junction 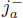 on *s*^−^ related via RC such that upstream and downstream are reversed:

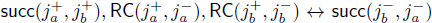

Further axioms can be added to state the relationship between base types on opposing strands (not shown here).

For any genome interval relation in *r* ∈ *R*^16^, we can define a reverse-complement cognate, *r*^*R*^. We obtain definitions for these automatically using the formula:

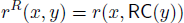

In table 4 we show only one `RC` relation, `upstream_overlaps`^*R*^. This is equivalent to `upstream_overlaps` with `RC` applied to the second argument. Note that unlike `upstream_overlaps`, this is a symmetric relation. Conversely, `nt_contained_by`^*R*^ (not shown), `is` the inverse of `nt_contains`^*R*^. See figure 2 for an illustration of why this is the case.

**Table 4:**
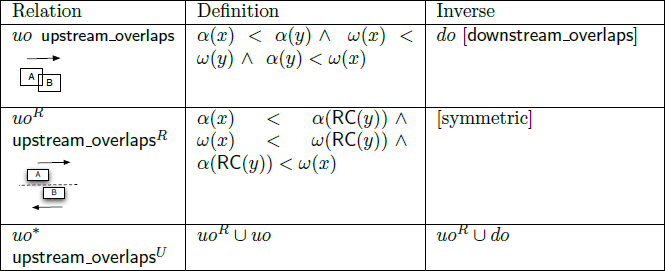
Example of Reverse Complementation cognate relations for the `upstream_overlaps` relation. Note the interaction between RC and inverse relations: in particular the inverse of *uo^R^* does not correspond to a single named relation.

| Relation | Definition | Inverse |
| --- | --- | --- |
| $uo$ **upstream_overlaps** | $\alpha(x) < \alpha(y) \wedge \omega(x) < \omega(y) \wedge \alpha(y) < \omega(x)$ | $do$ [downstream_overlaps] |
| $uo^R$ **upstream_overlaps**<sup>R</sup> | $\alpha(x) < \alpha(RC(y)) \wedge \omega(x) < \omega(RC(y)) \wedge \alpha(RC(y)) < \omega(x)$ | [symmetric] |
| $uo^*$ **upstream_overlaps**<sup>U</sup> | $uo^R \cup uo$ | $uo^R \cup do$ |

For any *r* ∈ *R*^16^, we can further define a relation *r^U^* = *r* ∪ *r^R^*. Table 4 illustrates this with `upstream_overlaps`^*R*^ ∪ `upstream_overlaps` which we call this `upstream_overlaps`^*U*^. These union relations correspond to common use cases (e.g. Region-of-Interest queries in Genome Browsers[10][11][12]), so we should have intuitive names for them. However, from the point of view of axiomatisation, it is simpler to treat these as derived rather than basic relations.

Thus we have 16 relations in the core set (including inverse relations for non-symmetric relations). We declare relations for this core set, plus their `RC` equivalents, plus the union set. This gives us 48 relations in total. This may seem excessive but as we will see we will need most for our use cases.

#### 2.1.4 Operations over collections of intervals

Many genomic features correspond to *collection* of intervals on a sequence - for example, the coding part of a multi-exon gene. Neither RCC-8 not AIA dictate operations over collections of regions or intervals. We propose a simple extension in GIA for dealing with such collections.

We overload the relations that are used for single intervals, but provide distinct definitions. We also define new relations that are only applicable to collections. For example, in figure 2 genes A and B stand in an *o^R^* (overlaps) relation, even though their respective exons share no bases. However, we can introduce a new relation `interleaves`^*R*^, defined such that the exons in gene A interleave (on the opposite strand) the exons in gene B. (note that it is important that we distinguish between the sets of exons interleaving and the genes overlapping).

We present here a subset of the full set of relations, which have yet to be finalized. We treat each collection as a set of intervals.

Overlaps must hold for some pair of elements from each pair:

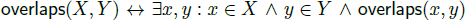

Adjacency must hold of any one pair, and in addition there should be no overlap:

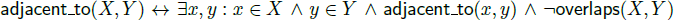

Upstream must hold for all elements:

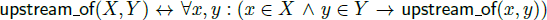

Interleaves is more complex:

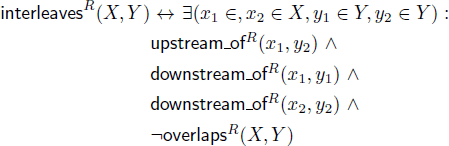

#### 2.1.5 Composition table

The full composition table for GIA(J) is too large to show but is available as part of the Genome Intervals relations file. Some examples include:

Transitivity of `upstream_of`:

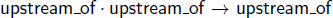

`starts` and `finishes` compose to make (non-tangential) containment: 
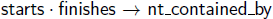

The RC upstream of relation is transitive over downstream of (consider Ax1 u Bx2 d Bx1 in figure 2):

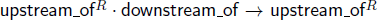

The full composition table was derived by an automated theorem prover (see methods).

### 2.2 Applications

### 2.3 Extending the Sequence Ontology

We can use the genome interval relations above to extend the SO, adding new logical definitions and constraint axioms – we call the resulting artefact `SO`^+^. These logical definitions can be used for both reasoning within the ontology, and to infer the presence of unstated genomic features in genome databases. The constraint axioms can be used to detect inconsistencies within the ontology, and to provide constraints for genome databases.

#### 2.3.1 Logical Definitions

Logical definitions provide necessary and sufficient conditions in computable form. We have created 140 new definitions for existing SO types based on the GIA relations. Table 5 shows a subset of these. SO uses the type-level version of these relations

**Table 5:** Examples of logical definitions of SO terms using GI relations. Relation quantifiers are taken to be All-Some unless otherwise noted. The semantics of a genus differentia definition (T,G,D) are such that *T*(*x*) ↔ *G*(*x*) ∧ ∀(*R*, *Y*) ∈ *D* : ∃*y*: *xRy*

| Type | Genus | Differentia |
| --- | --- | --- |
| five_prime_coding_exon | coding_exon | $\langle \text{overlaps start\_codon} \rangle$ |
| five_prime_intron | intron | $\langle \text{nt\_contained\_by five\_prime\_UTR} \rangle$ |
| UTR_intron | intron | $\langle \text{overlaps UTR} \rangle$ |
| twintron | intron | $\langle \text{nt\_contained\_by intron} \rangle$ |
| start_codon | codon | $\langle \text{starts CDS} \rangle$ |
| stop_codon | codon | $\langle \text{finishes CDS} \rangle$ |
| intergenic_region | region | $\langle \text{disconnected\_from gene} \rangle$<br>$\langle \text{upstream\_adjacent\_to gene} \rangle$<br>$\langle \text{downstream\_adjacent\_to gene} \rangle$ |
| noncoding_exon | exon | $\langle \text{disconnected\_from CDS} \rangle$ |
| intron | region | $\langle \text{nt\_contained\_by transcript} \rangle$<br>$\langle \text{upstream\_adjacent\_to exon} \rangle$<br>$\langle \text{downstream\_adjacent\_to exon} \rangle$<br>$\langle \text{disconnected\_from exon} \rangle$ |
| interior_intron | intron | $\langle \text{nt\_contained\_by CDS} \rangle$ |
| five_prime_UTR | mRNA_region | $\langle \text{upstream\_adjacent\_to CDS} \rangle$<br>$\langle \text{starts mRNA} \rangle$ |
| three_prime_UTR | mRNA_region | $\langle \text{downstream\_adjacent\_to CDS} \rangle$<br>$\langle \text{finishes mRNA} \rangle$ |
| interior_UTR | mRNA_region | $\langle \text{downstream\_adjacent\_to CDS} \rangle$<br>$\langle \text{upstream\_adjacent\_to CDS} \rangle$ |
| interior_exon | exon | $\langle \text{adjacent\_to five\_prime\_splice\_site} \rangle$<br>$\langle \text{adjacent\_to three\_prime\_splice\_site} \rangle$<br>$\langle \text{disconnected\_from splice\_site} \rangle$ |

Because the definitions are both necessary and sufficient, they can be used to construct queries on genome databases. For example, to find 5’ coding exons, make a conjunctive query for coding exons that overlap start codons. `overlaps` can be translated to a genome coordinate query. The query may need further expansion of start codon or coding exon is not materialized in the database.

#### 2.3.2 Relationship and Constraint Axioms

SO already has relationships between types specified in the ontology: for example, 〈`TSS` *part_of* `transcript`〉, which states that if *x* is an instance of a `TSS` then there must be some `transcript` *y* such that `part_of`(*x*, *y*).

Using the GIA relations we can add new relationships to SO, or refine existing relationships that may have unclear semantics. Table 6 shows a sample of these. Note that we exclude the relationships which are trivially obtained from the definitions in table 5.

**Table 6:** Proposed new relationship axioms for SO using genome interval relations. These are type-level relationships that can be translated to instance level relationships such that for example `TSS`(*x*) → ∃*y*, `primary_transcript`(*y*), `upstream_of`(*y*)

| Type | Relationship |
| --- | --- |
| TSS | $\langle \text{upstream\_of primary\_transcript} \rangle$ |
| polyA_sequence | $\langle \text{downstream\_adjacent\_to mRNA} \rangle$ |
| polyA_site | $\langle \text{end\_of mRNA} \rangle$ |

We can use these type-level relationships as constraints on a genome database. We must do this carefully, bearing in mind the fact that these axioms make an open-world assumption: just because an instance in reality is entailed to exist, it does not follow that this musy be explicitly represented in the genome database.

Note that many of the relationships in table 6 are in fact *underconstrained* as constraints. For example, take 〈`TSS upstream_of primary_transcript`〉. This is in fact an extremely weak axiom - it states that every `TSS` is upstream of *some* `primary_transcript` - the `TSS` and transcript do not have to be otherwise related. This will be trivially true: a `TSS` located at the end of a chromosome sequence will be upstream of *all* the other genes on the same strand.

In fact we want to say that every `TSS` lies upstream *of the same transcript that it regulates*. The current version of SO uses a *part_of* relation between the `TSS` and the `primary_transcript` to indicate that every `TSS` is functionally coupled to a `primary_transcript`. We can write a fully constrained axiom:

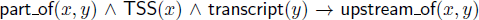

We can use this axioms to check instance-level data, as found in genome databases and data files.

### 2.4 Reasoning over genome intervals

We can use `SO`^+^(the combination of the SO extended with GIA relations) to perform reasoning tasks. These tasks can be broken down according to whether we are performing *validation* of axioms or *inference* of unstated axioms. We can further break this down according to whether we are performing reasoning over just `SO`^+^, or the combination of `SO`^+^ and some genomic instance data.

We can use a number of different systems for reasoning. Relational databases are generally considered to be the least expressive (meaning that we cannot translate some axioms to the relational model), but are also considered to scale to large genome-sized datasets. At the other extreme are first-order logic theorem provers, which allow for high expressivity, but do not scale well. In between there are a number of different systems that can be roughly divided into partially overlapping rule-based and description-logic (DL) approaches.

Rule based approaches extend the expressivity of the relational model with arbitrary recursive rules; the OBO-Edit reasoner is an example of a rule-based system. More expressive systems include disjunctive datalog engines such as DLV. OWL-DL (Web Ontology Language Description Logic) was designed to maximine expressivity and decidability, and there are a variety of OWL-DL reasoners.

The set of relations consisting of the core 16 basic GIA relations do not correspond to any of the tractable sub-algebras of the Allen Algebra (for example, the `adjacent_to` relation, defined in terms of Allen as *m* ∪ *mi* appears in none of the tractable subsets). However, this is not a major concern for the majority of genome databases in which interval junctions can always be assigned to discrete ordered bases.

We present first examples of reasoning using `SO`^+^, and then examples of reasoning using a combination of `SO`^+^ and genome databases.

#### 2.4.1 Reasoning over the ontology

Given an ontology consisting of a set of *asserted* relationships, a reasoner can infer the *entailed* relationships. The full set of entailed relationships for a relation *R* is called the *deductive closure* of *R*.

Computing entailed relationships is useful for ontology maintenance - the *is_a* hierarchy for 200 of the terms in SO is maintained automatically be the OBO-Edit reasoner, using the existing genus-differentia definitions.

Computing the deductive closure of all ontology relations is also useful for improving genome database queries.

Given that the SO is by itself several orders of magnitude smaller than a typical genome database, we can afford to use a system with higher expressivity to do the reasoning. We have used both OBO and OWL-DL reasoners to compute entailed relationships in `SO`^+^, and both give the same results.

In theory an OWL-DL reasoner can compute relationships that are difficult to compute using a rule-based approach. For example, inferring that every codon overlaps a CDS, based on the five axioms: (a) a codon is either a start or stop codon (b) start codons start a CDS, and (c) start implies overlaps (d) stop codons stop a CDS, and (e) stop implies overlap. Currently the OBO-Edit reasoner does not make use of class unions. In this particular case it doesn’t matter, as the relationship was already asserted at the codon level.

Figure 3 shows examples of entailed relationships. We can ask *how are the type 5’UTR and start codon related?* and get the answer `upstream_adjacent_to`, even though this fact is not explicitly stated in the ontology.

**Figure 3:**
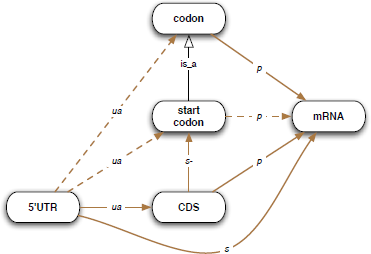
Reasoning over ontologies: this example illustrates the deductive closure involving a few SO types, and makes use of the relation composition axiom `upstream_adjacent_to · started_by` → `upstream_adjacent_to` to infer that every 〈`five_prime_UTR upstream_adjacent_to start_codon`〉 (inferred relationships are shown with dashed lines). Not all inferences are shown. This is an example of qualitative/symbolic reasoning - we can make inferences even without arithmetic, using the GIA

The other application of reasoning over ontologies is to find inconsistencies or unsatisfiable classes. Using both OBO-Edit and OWL-DL reasoners, we could find no inconsistencies between axioms within the extended SO. This is not surprising as the SO is carefully scrutinized by its editors prior to each release, and automated procedures are currently in use with the existing axioms. We still expect that the extended SO presents more precise axioms on which these reasoners can operate.

An example of a conceivable mistake that can be detected by reasoners is: (a) 〈`three_prime_UTR downstream_adjacent_to CDS`〉 (b) 〈`stop_codon` *is_a* `codon`〉 (c) 〈`codon nt_contained_by CDS`〉 (d) 〈`stop_codon starts three_prime_UTR`〉. This example is not entirely artificial: prior to the existence of SO there was inconsistency amongst the genomics community as to whether the stop codon should be considered part of the CDS. This inconsistency caused interoperability problems, which were solved for systems adopting SO.

## 3 Discussion

### 3.1 Enhanced rigor in the Sequence Ontology

In describing the GIA and extending SO we came across portions of SO that were in need of more precise textual definitions. For example, `intergenic_region` was defined as *A region containing or overlapping no genes that is bounded on either side by a gene*. In formalizing the definition for this using the `upstream_adjacent_to` and `downstream_adjacent_to` relations to represent *bounded on either side* we realized that this definition excluded the two regions on either end of a chromosome. The textual definition was extended to be include the disjunctive clause *or bounded by either a gene or the end of the chromosome*. Another example was `splice_site`, which had a textual definition indicating that it was a junction but a placement in the *is_a* hierarchy indication a region. Once the computable definition was added, the inconsistency could be detected by a reasoner (although it was in fact detected whilst preparing the logical definition). This resulted in the definition being clarified and the addition of a new term splice_junction.

### 3.2 Junction-oriented vs Base-oriented

Our formulation is a junction or *interbase* one. We could equally have defined interval relations in terms of the bases themselves. We consider an interbase system to have a slight advantage in terms of simplicity of representation of positioning of splice junctions, insertion regions and so on. However, the two systems would be equivalent in terms of expressivity, so the choice of one over the other is arbitrary.

### 3.3 Unresolved Issues

#### 3.3.1 Splicing, transcripts and exon identity

The examples presented in this paper make a simplifying assumption, namely that all types in the SO represent features along a DNA sequence. The relationships and definitions presented here do not account for the biology of splicing and translation. For example, exons are non-adjacent on the DNA sequence and on the unspliced RNA sequence, but become adjacent after splicing. A full treatment will have to account for this temporal aspect. One approach is to introduce different types, such as exon^*G*^ and exon^*T*^ for exons on genomes and exons on transcripts respectively. Another is to use n-ary relations such that it is possible to state 〈`exon adjacent_to exon` *on* `mature_transcript`〉 and 〈`exon disconnected_from exon` *on* `genome`〉.

#### 3.3.2 Circular genomes

The current axiomatization needs to be modified to handle circular genomes. One approach is to weaken the definition of < such that it is reflexive on circular genomes. However, this will have some consequences for the other axioms, which needs to be fully worked out. For example, every feature would be upstream of every other feature.

Another solution would be to have some kind of probabilistic metric or arbitrary cut-off, whereby junctions are no longer considered upstream if they loop around the circular DNA too far. But this would be difficult to integrate into these existing axioms.

The most likely approach is to use origin of replication (oriC in bacteria) as the origin of the sequence.

## 4 Conclusions

We have defined a collection of genome interval relations, and used them to define an extension of the Sequence Ontology (SO). This extension and these relations will soon become part of the core SO. The relations help clarify the meaning of terms in the SO to humans, and can be used by automated reasoning systems to assist with the construction and quality control of the ontology.

In additions the relations are useful for querying over and checking the conformance of datasets to constraints in the SO. The relations help clarify the meaning of certain queries to human beings, such that a query for “all exons upstream of gene ABC” has precise semantics. The extended SO can be used to enhance database queries such that implicit features such as introns are found. We found the most effective system for querying over genome datasets was a relational database with the help of a query expansion system. OWL-DL systems do not yet scale over genome sized datasets.

The composition table is useful for making inferences over the ontology, but for making inferences over data it is simpler to use arithmetic definitions rather than a composition table.

We believe that as genome datasets grow in size, complexity and importance the need for formal computable specifications of genomic data will increase. Specifications such as the one outlined here will be vital for ensuring the semantic and biological correctness of important datasets, and for performing advanced queries over these datasets.

## 5 Methods

### 5.1 Defining the Genome Interval Relations

We used OBO-Edit[13] to specify the genome interval relations and to create an extended subset of SO that used these relations for genus-differentia definitions. This was stored in an OBO Format 1.3 file.

### 5.2 Generation of composition table

To generate the full composition table, we first translated the genome interval relations to Prover9 syntax and used the Prover9 tool to calculate the table by brute force attempts to prove every possible *R*1 · *R*2 → *R* for all values of *R*, *R*1 and *R*2.

## 6 Availability

Common Logic specifications of the relations can be obtained from the github repository ^2^

## Footnotes

1 sometimes called “during”

2 https://github.com/cmungall/genomize-intervals-formalization

## References

1 B. Smith, W. Ceusters, J. Kohler, A. Kumar, J. Lomax, C.J. Mungall, F. Neuhaus, A. Rector, and C. Rosse. Relations in Biomedical Ontologies. Genome Biology, 6(5), 2005. URL http://genomebiology.com/2005/6/5/R46.

2 K. Eilbeck, S. E. Lewis, C. J. Mungall, M. D. Yandell, L. D. Stein, R. Durbin, and M. Ashburner. The Sequence Ontology: a tool for the unification of genome annotations. Genome Biology, 6(5), 2005.

3 P. Grenon and B. Smith. SNAP and SPAN: towards dynamic spatial ontology. Spatial Cognition & Computation,4(1):69–104, 2004.

4 E. Birney, T. D. Andrews, P. Bevan, M. Caccamo, Y. Chen, L. Clarke, G. Coates, J. Cuff, V. Curwen, T. Cutts, T. Down, E. Eyras, X. M. Fernandez-Suarez, P. Gane, B. Gibbins, J. Gilbert, M. Hammond, H. R. Hotz, V. Iyer, K. Jekosch, A. Kahari, A. Kasprzyk, D. Keefe, S. Keenan, H. Lehvaslaiho, G. McVicker, C. Melsopp, P. Meidl, E. Mongin, R. Pettett, S. Potter, G. Proctor, M. Rae, S. Searle, G. Slater, D. Smedley, J. Smith, W. Spooner, A. Stabenau, J. Stalker, R. Storey, A. Ureta-Vidal, K. C. Woodwark, G. Cameron, R. Durbin, A. Cox, T. Hubbard, and M. Clamp. An overview of Ensembl. Genome Res, 14(5):925–8, 2004. 1088-9051 Journal Article Review Review, Tutorial.

5 Christopher J. Mungall, David B. Emmert, and The FlyBase Consortium. A Chado case study: an ontology-based modular schema for representing genome-associated biological information. Bioinformatics, 23(13):i337–346, 2007. doi: 10.1093/bioinformatics/btm189. URL http://bioinformatics.oxfordjournals.org/cgi/content/abstract/23/13/i337.

6 Barry Smith and Anand Kumar. Controlled vocabularies in bioinformatics: a case study in the gene ontology. Drug Discovery Today: BIOSILICO, 2(6):246–252, 2004. TY - JOUR.

7 A G Cohn, B Bennett, J Gooday, and N Gotts. Representing and Reasoning with Qualitative Spatial Relations about Regions. In O Stock, editor, Temporal and spatial reasoning. Kluwer, 1997.

8 J. F. Allen. Maintaining knowledge about temporal intervals. Communications of the ACM, 26:832–843, 1983.

9 Andrei Krokhin, Peter Jeavons, and Peter Jonsson. Reasoning about temporal relations: The tractable subalgebras of Allens interval algebra. Journal of the ACM, 50:2003, 2003.

10 L. D. Stein, C. Mungall, S. Shu, M. Caudy, M. Mangone, A. Day, E. Nickerson, J. E. Stajich, T. W. Harris, A. Arva, and S. Lewis. The generic genome browser: a building block for a model organism system database. Genome Res, 12(10):1599–610, 2002.

11 M.E. Skinner, A.V. Uzilov, L.D. Stein, C.J. Mungall, and I.H. Holmes. JBrowse: A next-generation genome browser. Genome Research, 2009. URL http://genome.cshlp.org/content/19/9/1630.long.

12 S. E. Lewis, S. M. Searle, N. Harris, M. Gibson, V. Lyer, J. Richter, C. Wiel, L. Bayraktaroglir, E. Birney, M. A. Crosby, J. S. Kaminker, B. B. Matthews, S. E. Prochnik, C. D. Smithy, J. L. Tupy, G. M. Rubin, S. Misra, C. J. Mungall, and M. E. Clamp. Apollo: a sequence annotation editor. Genome Biol, 3(12):0082–1, 2002.

13 John Day-Richter, Midori A Harris, Melissa Haendel, Gene Ontology OBO-Edit Working Group, and Suzanna Lewis. OBO-Edit–an ontology editor for biologists. Bioinformatics, 23(16):2198–2200, Aug 2007, doi: 10.1093/bioinformatics/btm112. URL http://dx.doi.org/10.1093/bioinformatics/btm112.

